# Expanded Proteome Coverage Powered by Advanced Ion Processing Enables Deep Single-Cell Drug Response Subtyping in Human Stem Cell Derived Cardiomyocytes

**DOI:** 10.64898/2026.05.29.728818

**Authors:** Johannes V Janssens, Aleksandra Binek, Lizhuo Ai, Ajay Bharadwaj, Madelyn Arzt, Diego Assis, Matthew Willetts, Michael Krawitzky, Daniel Hornburg, Arun Sharma, Aleksandr Stotland, Jennifer E. Van Eyk

**Author notes:** **Corresponding Authors:** Dr Jennifer E Van Eyk, Professor, Smidt Heart Institute, Director, Advanced Clinical Biosystems Research Institute, Erika Glazer Endowed Chair in Women’s Heart Health, Cedars-Sinai Medical Center, Los Angeles, CA, Dr Aleksandra Binek, Project Scientist, Smidt Heart Institute, Advanced Clinical Biosystems Research Institute, Proteomics and Metabolomics Core, Cedars-Sinai Medical Center, Los Angeles, CA. Authors contributed equally (Johannes V Janssens, Aleksandra Binek). Authors contributed equally (Lizhuo Ai, Ajay Bharadwaj). Authors contributed equally (Aleksandr Stotland, Jennifer E. Van Eyk).

## Abstract

Single-cell proteomics (SCP) enables the study of cellular heterogeneity at the functional level but remains limited by incomplete proteome coverage and high data missingness. Here, we present an enhanced label-free SCP workflow that leverages the timsUltra AIP mass spectrometry platform equipped with the Athena Ion Processor (AIP). Across a controlled dilution series of human induced pluripotent stem cell-derived cardiomyocytes (iPSC-CMs), AIP-enabled acquisition consistently increased proteome depth and detection consistency across cells at all input levels. In single iPSC-CMs, the timsUltra AIP quantified up to 3,858 protein groups, averaging ∼1,300 proteins per cell, enabling robust proteome-level classification of cardiomyocyte subtypes. Using a reference-based protein classifier, cells were stratified into mature cardiomyocytes and less differentiated cell states, revealing substantial baseline heterogeneity. Importantly, increased single-cell sensitivity translated directly into biological insight, as approximately 30% of differentially expressed proteins associated with subtype-specific drug responses were detected exclusively by timsUltra AIP. Application of this workflow to PR-364 (a mitophagy boosting drug) dose-response experiment uncovered distinct, subtype-dependent pathway adaptations. Mature cardiomyocytes exhibited dose-dependent increases in mitochondrial and metabolic pathway activity, while immature cells showed enrichment of cytoskeletal and developmental programs. These effects were partially obscured in simulated bulk analyses, highlighting the value of single-cell resolution. Together, these results demonstrate that improved fragment ion transmission and utilization translate directly into enhanced biological insight, enabling more comprehensive and functionally relevant single-cell proteomics.

## Introduction

Single-Cell Proteomics (SCP) has emerged as a powerful approach for resolving cellular heterogeneity at a functional level, complementing transcriptomic measurements by directly capturing protein abundance and pathway activity at single-cell input levels. However, SCP remains constrained by limited sensitivity, incomplete proteome coverage, and high levels of data missingness, particularly for low-abundance proteins. These challenges are especially limiting when resolving subtle biological differences, such as cell state transitions[1], inference of temporal progression (pseudo-time analysis)[2], cell fate trajectories, lineage heterogeneity[3, 4], or drug-induced responses[5].

Advances in mass spectrometry instrumentation[6-8] and acquisition schemes (data-independent acquisition, DIA) have improved peptide identification rates and quantitative consistency, yet efficient utilization of fragmented ion information remains a bottleneck in isolated single-cell proteomics. Enhancing the transmission and detection of peptide fragment ions without introducing bias could increase consistent proteome depth while preserving quantitative accuracy. The recently introduced timsUltra AIP deploys a novel Athena Ion Processor (AIP) in conjunction with front end trapped Ion Mobility Separation (TIMS). AIP has been engineered to significantly increase fragment ions transmission efficiency from the collision cell to the orthogonal accelerator of the time-of-flight mass analyzer. Since AIP has been integrated into timsTOF Ultra 2 framework, we hypothesize it to be particularly impactful for high sensitivity applications such as single cell proteomics. We benchmarked timsUltra AIP performance across controlled low quantity iPSC-CM dilution series and then applied the workflow to characterize iPSC-CM subtype heterogeneity and drug-induced proteomic remodeling at single-cell resolution. Improved peptide fragment ion transmission in the new timsUltra AIP in combination with trapped ion mobility and data independent acquisition (DIA) facilitated an increase in proteome depth, improved data completeness, and enabled more reliable quantification of single iPSC-CMs.

Human iPSC-cardiomyocytes are a well-established translational *in vitro* model system. iPSC-CMs bridge research in cardiac biology, disease mechanisms, and pharmacological responses carried out in animal models[9] and as part of phase 1 clinical trials of drug candidates[9]. Importantly, however, iPSC-CMs display intrinsic cellular heterogeneity[4], encompassing a spectrum of maturity phenotypes with distinct metabolic and contractile structural characteristics. Specifically, we found differentially expressed proteins that revealed cell clusters expressing cardiac-specific proteins (*e.g*. myosin heavy chain 6, MYH6) and cell clusters with significantly higher smooth muscle cell protein markers, including Caldesmon (CALD1), Myosin-11 (MYH11), Myosin light chain 6 (MYL6), and Myosin light chain 12B (MYL12B)[4]. Capturing this heterogeneity at the single-cell proteome level requires both high sensitivity and robust quantification. As proof-of-principle, we used the previously described small molecule PR-364[9], which activates Parkin, an E3 ubiquitin ligase. Activation of Parkin increases clearance of dysfunctional mitochondria and maintains cardiomyocyte mitochondrial homeostasis, resulting in improved heart function following a myocardial infarct[9]. Here, we evaluate the heterogeneous response of human iPSC-CM subtypes to PR-364 in culture using single-cell proteomics, focusing on changes in central metabolomic pathways and protein expression differences in human iPSC-CMs as part of a translational step towards clinical application.

## Results

### Enhanced single cell proteome depth and quantitative performance with application of AIP

We first assessed the impact of AIP on protein identification and quantitative robustness across a dilution series of control iPSC-CMs. 1, 3, 5, 10, or 15 cells were dispensed per well into a 384-well PCR plate. Samples were processed using an identical workflow and liquid chromatography (LC) settings, followed by mass spectrometry (MS) analysis with and without AIP (**Fig.1A**). On the timsUltra AIP, the number of proteins identified per sample across the entire cell dilution series were consistently increased. The most pronounced gain was observed at the lowest quantities. The single cell samples exhibited a more than two-fold increase in protein identifications compared to conventional ion processing (No AIP: 409 compared with AIP: 1039, n=48-50 cells per group, p<0.05). The distribution of protein counts across cell quantities further highlighted the most significant improvements at single cell and lowest cell dilutions (**Fig. 1B**) and in iPSC-CMs protein lysate dilutions (5ng, 1ng, 500pg, 100pg, **Fig. S1A-B**). Similar missed cleavage rates between AIP and No AIP confirm similar digestion characteristics for iPSC-CMs in each group (**Fig. 1C**). Quantitative reproducibility in AIP datasets showed modest improvement with a consistent shift toward lower percent coefficient of variation (%CV) values, and higher proportion of proteins falling within the <20% CV bin (**Fig.1D)**. Precision of timsUltra AIP in protein lysate samples yielded over 75% of proteins with %CV values below 20% at the higher loads (5 ng and 1 ng) (**Fig. S1C–D)**. Missingness patterns across 1797 commonly detected proteins within both acquisition schemes revealed a noticeable reduction in missing values in single cell dataset acquired with AIP (**Fig. 1E**). Importantly, the observed detection improvements were not driven by biases in peptide physicochemical properties, as the distributions of peptide lengths and charge states were closely overlapping (**Fig1. F-G**), suggesting that enhanced ion transmission efficiency does not alter/favor any peptide population.

**Figure 1.**
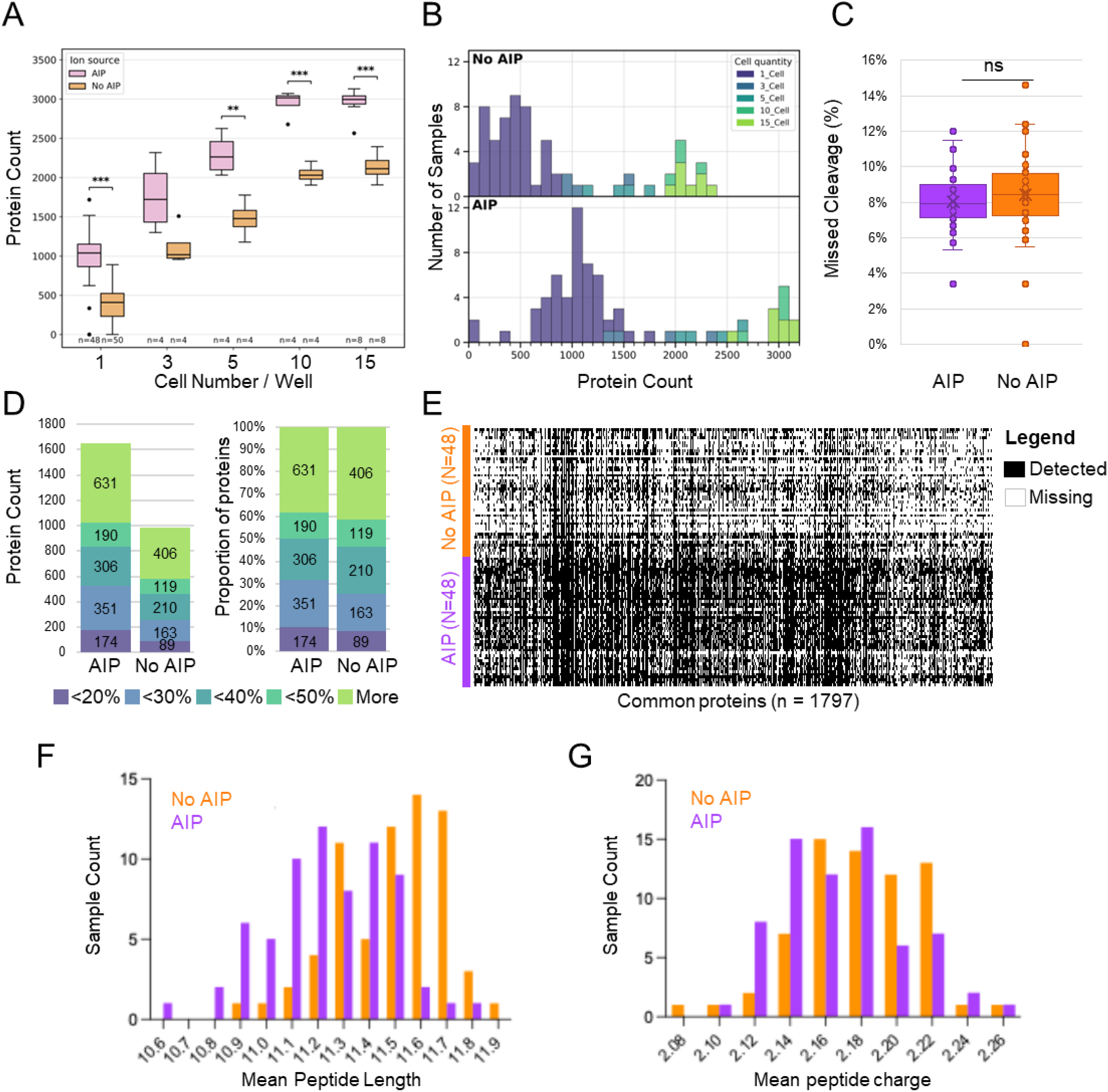
timsUltra AIP improves protein identification and reduces data missingness in low-input samples when compared to timsTOF Ultra2. **A)** Total protein counts identified per sample across increasing cell inputs of iPSC-CMs (per 1, 3, 5, 10, and 15 cells) processed with the timsUltra AIP or timsTOF Ultra 2. Boxplots show median and interquartile range, and whiskers indicate variability across replicates. We detected statistical significance across all conditions of the cell serial dilution curve (*p < 0.05; **p < 0.01; ***p < 0.001). **B)** Distribution of protein counts per sample for datasets generated on timsTOF Ultra 2 (top) and timsUltra AIP (bottom). Colors indicate different amounts of cell input, demonstrating increased depth and consistency of protein identifications with timsUltra AIP. **C)** Percentage of missing values per sample for timsUltra AIP and regular collision. Boxplots show missingness across samples (ns indicates no significant difference). **D)** Number of protein identifications stratified by percent coefficient of variation (%CV) for both systems. The left panel shows absolute counts, and the right panel shows percentages. **E)** Missingness heatmap of proteins commonly detected across samples from both conditions (n = 1,797 proteins). Black indicates observed values and white indicates missing values. Samples are grouped by system configuration (AIP vs no AIP). **F)** Distribution of average peptide lengths identified. **G)** Distribution of average peptide charge states identified.

### Detection of low-abundance proteins expands functional proteome coverage

Proteome composition was evaluated by comparing protein overlap and abundance distributions across all single-cells for the timsTOF Ultra 2 and timsUltra AIP (**Fig. 2A-C**). This revealed that a substantial proportion of proteins (n=1034), over half of the total identified proteome was uniquely identified with the new timsUltra AIP while only 6 proteins were exclusive to the previous generation system. Ranked abundance plots across both datasets show that AIP provides a clear advantage in the detection of low-abundant proteins (**Fig. 2B**). Most mean protein abundance values aligned closely along the diagonal, indicating strong agreement between both systems in terms of quantification for shared proteins (**Fig.2C**). However, a noticeable subset of proteins deviated above the lower part of the diagonal, suggesting increased signal intensity under AIP conditions, particularly for lower-abundance proteins. We classified all detected proteins from single cell samples based on abundance: detected only with timsUltra AIP (red), statistically higher with timsUltra AIP (blue), similar in both (green), and statistically lower with timsUltra AIP (grey). This analysis showed that a quarter of the shared proteome exhibited higher intensities in the AIP dataset, while a very small fraction showed higher intensity in the timsTOF Ultra 2 data (**Fig. 2D**). Importantly, a substantial subset of proteins (∼50%) was uniquely detected with the timsUltra AIP. The inset pie chart highlights the distribution of these categories, illustrating that the AIP contributes to both-increased detection and enhanced signal for a large portion of the proteome. As expected, Gene Ontology Biological Processes (GO: BP) pathway coverage terms (**Fig. 2E**), consistently increased with the higher sensitivity timsUltra AIP, as reflected by larger dot sizes across the entire range of biological processes. Similarly, GO Cellular Component (GO: CC) analysis (**Fig.2F**) showed expanded coverage of proteins associated with subcellular structures and complexes, including mitochondria, cytoskeletal elements, and membrane-associated compartments.

**Figure 2.**
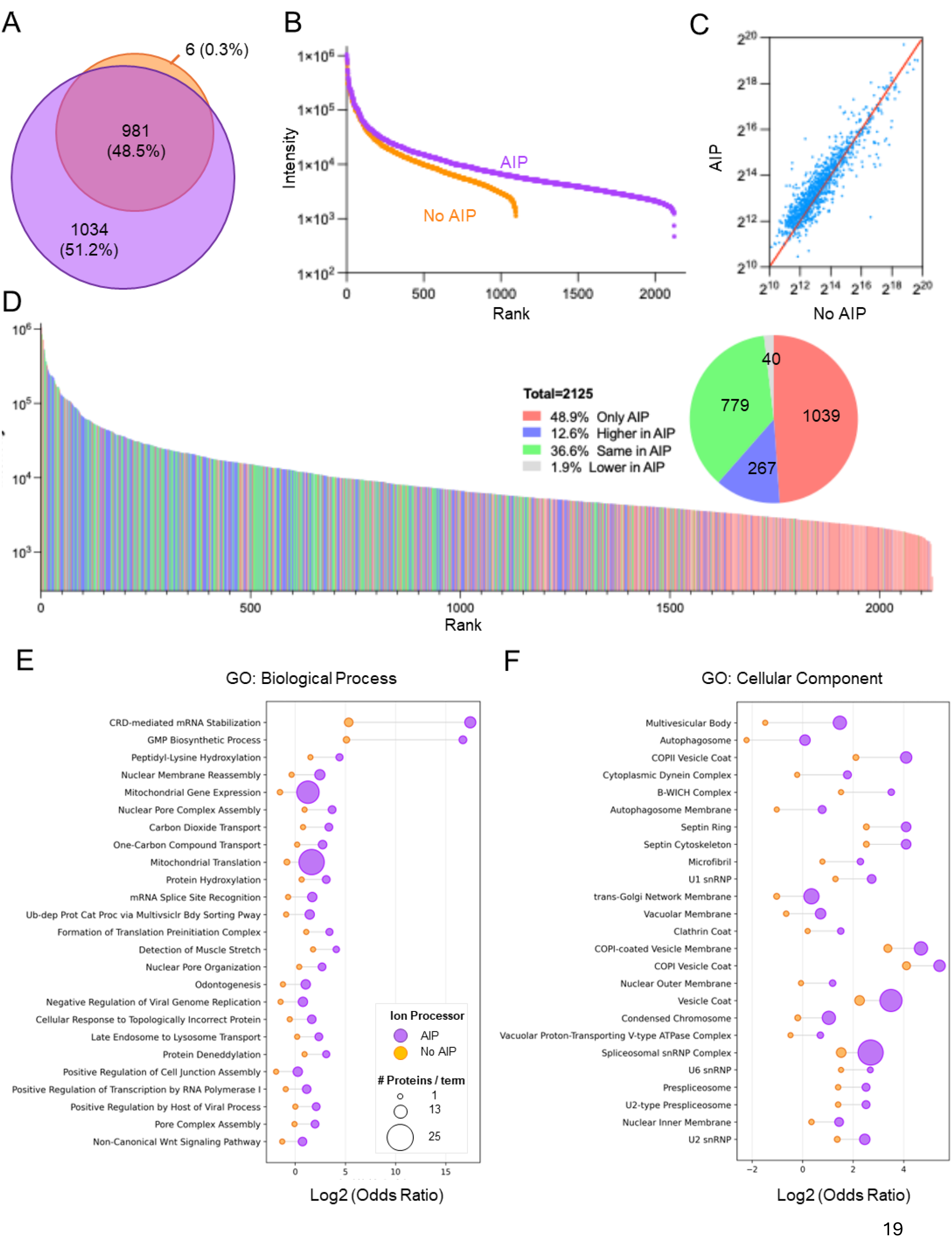
Impact of timsUltra AIP on protein overlap, abundance, and GO terms and pathways coverage. **A)** Venn diagram showing overlap of proteins identified with the timsUltra AIP vs. timsTOF Ultra 2. Percentages indicate the proportion of total identified proteins. This comparison shows only proteins identified in at least 10% of the samples in each dataset. **B)** Ranked abundance plot of all identified proteins across two datasets, comparing protein signal intensities between AIP and no-AIP conditions. Proteins are ranked in decreasing abundance, showing that AIP performs better on low-abundant proteins. **C)** Scatter plot comparing mean protein abundances between no-AIP and AIP datasets. Each point represents a protein; the diagonal line indicates equal abundance. The cloud of protein points above the diagonal line suggests the improved performance of AIP in lower-abundance proteins. **D)** Ranked abundance plot for all AIP proteins. Proteins are colored according to whether they show higher abundance in AIP (blue), higher abundance in no AIP (grey), similar abundance (green), or presence in AIP only (red). The inset pie chart summarizes the proportion of proteins in each category (total = 2,125 proteins). No proteins were filtered out based on their presence in either dataset. **E)** Gene Ontology Biological Process (GO: BP) pathways showing increased protein coverage with AIP. Dot size reflects the number of proteins mapped to each pathway, and color indicates the ion processing condition. **F)** Gene Ontology Cellular Component (GO: CC) pathways showing increased protein coverage with AIP, displayed as in panel E.

### High-resolution mapping of iPSC subtypes and drug response heterogeneity

Next, we sought to investigate the utility of the timsUltra AIP for making biological discoveries in a high sensitivity single-cell proteomics experiment. To this end, iPSC-CMs were subjected to a dose-response treatment with the selective parkin activator PR-364. iPSC-CMs were analyzed at four concentrations (0 µM, 3 µM, 10 µM, and 20 µM, **Fig. 3A**), and proteomic profiles were projected into a low-dimensional space by performing uniform manifold approximation and projection (UMAP) analysis (**Fig. 3B**). The resulting protein abundance data embeddings revealed no clear global separation of cell populations across drug concentrations, or their alignment with Leiden clustering, which delineated 8 distinct cluster subpopulations. Nevertheless, marked heterogeneity of iPSC-CM maturity markers MYH6 and MYH7 was observed (**Fig.S2A**) leading us to hypothesize that the iPSC-CMs incorporated in the study may exist at varying stages of differentiation. Using protein markers we have previously linked with iPSC-CM differentiation status as classifiers, we embarked on an approach to annotate iPSC-CMs as ‘mature cardiomyocytes (CMs)’ or ‘immature cells’[4]. ‘Immature cells’ for the purpose of subtype naming in the current study were not neonatal or fetal forms of cardiomyocyte. They were simply cells that exhibited a proteome that had not fully differentiated into cardiomyocytes. They expressed proteins commonly associated with other cell types or the developing heart, but not the adult heart. Briefly, iPSC-CMs were classified into 2 subtypes (**Fig.3B, *top right***) using the calculated difference between the mean of MYH7 + MYH6 and the mean of TPM4 + FLNA expression (see methods for further detail) which we termed ‘mature CMs’ (Leiden clusters 0, 1, 5) and ‘immature cells’ (clusters 2, 3, 4, 6, 7). 44% (N = 124 / 284) iPSC-CMs were classified into the mature category. iPSC-CMs defined as ‘mature CMs’ by the binary maturity classifier outlined above exhibited higher expression of Myosin heavy chain 6 (MYH6), as expected. Reassuringly, iPSC-CMs designated as ‘immature cells’ showed elevated levels of smooth muscle (i.e. non-cardiomyocyte) proteins Caldesmon 1 (CALD1), Myosin light polypeptide 6 (MYL6), and Myosin regulatory light chain 12A (MYL12A) (**Fig. 3B & S2B**) which were not included in the classifier function. Composite maturity and immaturity scores as well as maturity ratio including the smooth muscle proteins provided additional support for cellular subtype separation (**Fig.3B, S2A&B**). Key DEPs were selected and overlayed onto the UMAP embedding, including Ubiquitin carboxy-terminal hydrolase L1 (UCHL1), FK506-Binding Protein 1A (FKBP1A), Transgelin (TAGLN), and Vimentin (VIM). Those projections displayed distinct spatial patterns aligned with the maturity scoring annotation and captured both known and novel markers of iPSC-CM heterogeneity.

**Figure 3.**
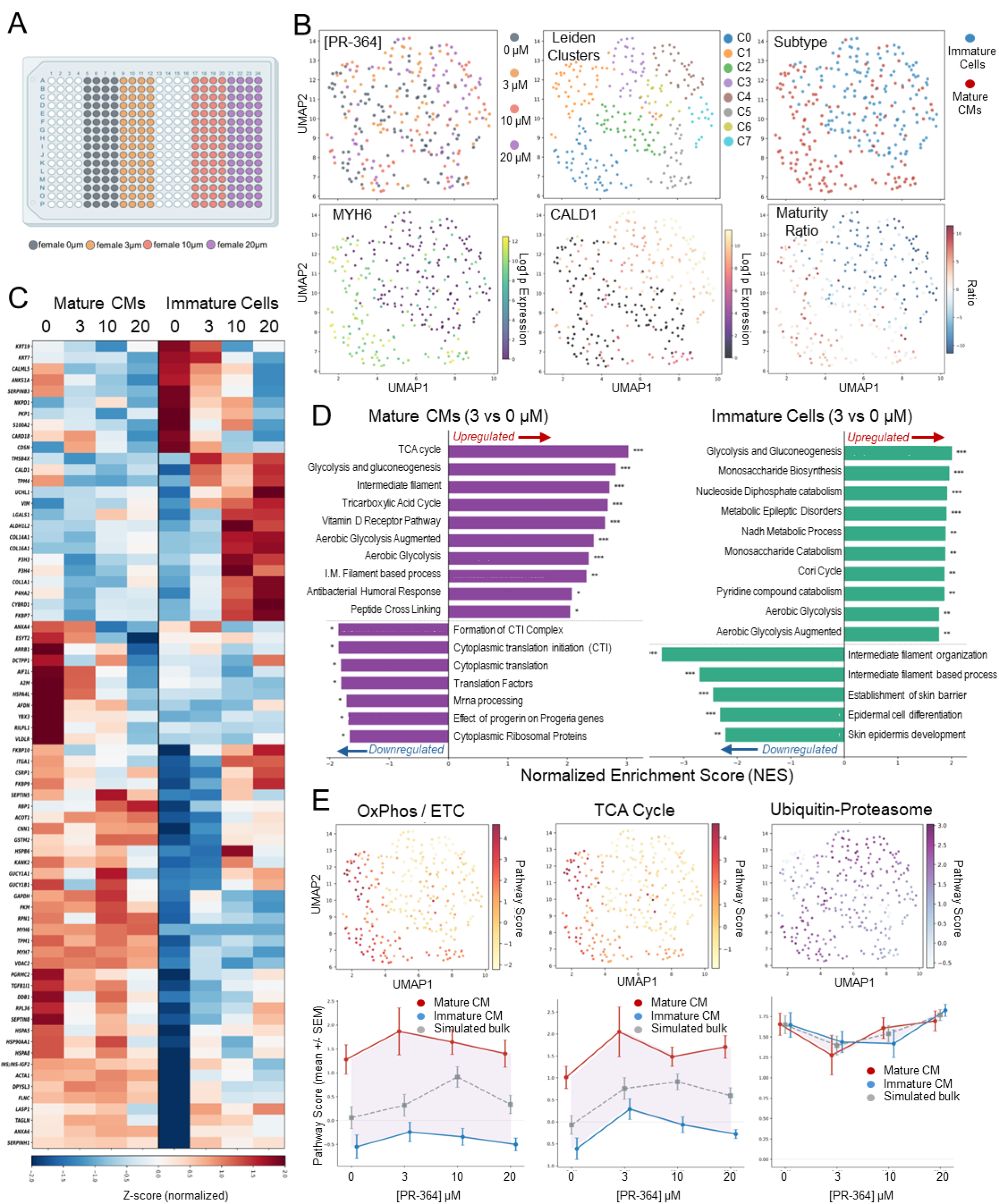
timsUltra AIP -enhanced single-cell proteomics reveals improved depth, clustering, and functional resolution across iPSC-CMs subtypes. **A)** Experimental layout of a 384-well plate with female-derived isolated single cell iPSC-CMs samples in PR-364 drug dose response (0 µm, 3 µm, 10 µm, and 20 µm). **B)** UMAP projections of single-cell proteomes under different drug concentrations. Left: baseline (0 µm) and 3 concentrations of PR-364. Middle: Leiden clustering highlighting distinct cell populations. Right: CMs subtype annotation (reference protein-based classification). Lower panels show protein intensity overlays for representative markers, including MYH7 (mature cardiomyocyte marker), CALD1 (immature marker), and computed maturity scores, illustrating improved biological separation with AIP. **C)** Protein identification counts stratified by drug dose and iPSC-CMs subtypes. Boxplots demonstrate increased protein detection with AIP across all groups, with the most pronounced gains at lower sample amounts. **D)** Heatmap of top 10 differentially expressed proteins (adjusted p-value < 0.01) across each drug concentration group in mature and immature iPSC-CMs clusters. Hierarchical clustering reveals clear subtype-specific proteomic signatures and enhanced contrast between the groups. **E)** Pathway enrichment analysis of differentially expressed proteins for mature (left) and immature (right) iPSC-CMs populations at 3 µm PR-364. Bar plots show significantly enriched biological processes, with green indicating GO Biological Process pathways and red indicating WikiPathways. **F)** Functional per-cell activity inference in drug dose-response. Top panels: UMAP overlays of inferred metabolic pathway activities, including oxidative phosphorylation/electron transport chain, TCA cycle, and ubiquitin–proteasome system. Bottom panels: dose-response curves (single-cell vs. simulated bulk) for female iPSC-CMs, demonstrating differential trends within the mature and immature cell subtypes with improved sensitivity enabled by AIP measurements.

### Pathway activity analysis reveals subtype-specific and dose-dependent functional responses to drug treatment

Distinct proteomic signatures associated with drug response were identified for each cell subtype (**Fig.3C**). Hierarchical clustering of the top10 differentially expressed proteins for 0 vs 3uM, 0 vs 10uM and 0 vs 20uM of PR-364 within each subtype revealed clear separation between mature CMs and immature cells across all drug concentrations. Crucially, approximately 30% of DEPs between subtypes and conditions were detected exclusively with the timsUltra AIP underscoring the relevance of highest sensitivity to monitor biological changes in single cell setups (**Supplementary Table** S**1**). Pairwise pathway enrichment analysis was carried out on DEPs to gain insight into the biological implications of PR-364 treatment. At 3 µM (relative to 0 µM), mature CMs showed upregulation of proteins enriched in pathways related to energy metabolism and contractile function, while immature cells were downregulated in pathways associated with cellular filaments organization (**Fig.3D**). At all doses, immature cells consistently exhibited the same trends in enrichment of developmental and structural pathways, whereas mature CMs showed upregulated enrichment of metabolic and mitochondrial processes (**Fig. S3**). Notably, there were instances in which increasing drug concentrations were associated with stronger pathway enrichment reflected in gradually and significantly increasing normalized enrichment scores (NES). Additionally, we inferred per-cell pathway activity scores and projected them onto UMAP embeddings (**Fig. 3F, Fig.S4**). This approach revealed 2-dimensional spatial gradients of pathway per-cell activity corresponding to distinct iPSC-CMs subtypes. OxPhos and TCA cycle activity were enriched in mature cardiomyocytes, consistent with their higher metabolic demands[10]. Conversely, pathways related to protein homeostasis, and cytoskeletal organization showed distinct distributions across subtypes, reflecting functional specialization (**Fig.3E, top panels, and Fig.S4C**). Dose-response analysis of pathway activities revealed distinct trends between mature CMs and immature cells (**Fig. 3E, bottom panels**). For example, metabolic pathways showed increased enrichment with higher drug concentrations in mature CMs, while immature cells exhibited more variable responses. Comparison with simulated bulk profiles demonstrated that single-cell measurements capture additional layers of heterogeneity that are obscured in bulk analyses. In **Supplementary Figure S4**, (panels A-B, C-D) we examined additional pathways, including collagen biosynthesis, wound healing, glycolysis, and pyruvate metabolism, which revealed complex, pathway-specific responses to drug treatment, including non-linear dose-response relationships and iPSC-CM subtype-specific dynamics.

## Discussion

We report the first application of Athena Ion Processor (AIP) as part of the timsTOF Ultra 2 framework for single-cell proteome analysis of human iPSC-CMs. We establish the analytical performance of the timsUltra AIP enhanced SCP workflow and provide a case example, highlighting how expanded deep proteome profiling and quantification can improve pathway analysis and reveal the heterogeneity of differentiation and variable drug response at the single cell proteome level.

For SCP to progress as a field improved recovery of low-abundance biology is essential. Smarter acquisition methods that reduce co-isolation, implementation of improved fragment ion utilization, and computational strategies that extract quantitative spectral signal from very low ion counts are example approaches. In SCP, data matrices are sparse and intensity dependent [11-14]. Missing SCP data points between single cell acquisitions can represent lack of sampling or technical variation rather than the protein absence in the specimen. Cardiovascular cells add challenges, because of small size and dramatically varying sarcomeric/mitochondrial content across cell types, within a particular cell type (*e.g*., cardiomyocytes) and across disease states [15-18], emphasizing that the method of MS acquisition needs to be optimal and consistent.

In the present work, we show the timsUltra AIP substantially enhances low abundance protein detection and quantification performance in iPSC-CMs, enabling deeper proteome coverage, reduced data missingness, and maintains robust quantitative accuracy without analytical bias (based on missed cleavage or peptide physicochemical properties). Our data indicates that these improvements substantially expand proteome coverage rather than simply redistributing the intensity signal across already detected proteins. Our key finding, especially at the single-cell level, shows a more than two-fold protein detection improvement with maintained quantitative reproducibility (lower coefficients of variation, with more proteins quantified, CV <20%). In studies with subtle biological perturbations large percentage of missing values can conceal meaningful signals. Data missingness across many individual cells, which is a persistent challenge in SCP, was reduced across commonly detected proteins. Improvement in transmission of informative fragment ions seems to be particularly advantageous for low-abundant peptides. Strikingly, approximately half of the detected proteome was uniquely identified with the timsUltra AIP, underscoring its ability to expand the proteomic landscape. Importantly, that expanded proteome contributed substantially to increased GO pathway coverage as illustrated in **Figure 2**. Thus, AIP not only increases the number of detected proteins but also enhances the detection of biologically relevant, low-abundant proteins that enable more comprehensive functional analysis.

Human samples are necessary in the pre-clinical cardiology field, with appreciation of the limitations of working with rodent tissues and cells[19, 20]. Even in clinical centers and hospitals, human cardiac biopsies can be scarce. Human iPSC-CMs, which can be differentiated from readily available sample types (blood, skin), represent a powerful and more available pre-clinical cardiac preparation amenable to genetic manipulation and drug treatment. iPSC-CMs are known to exhibit an immature-like phenotype characterized by spontaneous beating, disorganized sarcomere structures, and dependence on anaerobic metabolism[21-24]. However, they possess functional cardiac ion channels, respond to drug treatment, and can model patient-specific genetic backgrounds, despite their structural immaturity[22, 24-27]. Inherent to iPSC-CM preparations is a large degree of heterogeneity, which has been previously observed in functional[28] and transcriptomic evaluations[29] and recently the proteome[4] even at single differentiation timepoints. While standardization of differentiation approaches is an ongoing endeavor in the field, we have recently used single-cell proteomics to identify markers of iPSC-CM differentiation stage[4].

We have also incorporated some of the protein markers identified in Ai *et. al*.[4] into a binary classification system to derive two iPSC-CM subtypes likely linked to differentiation stage. While global clustering did not segregate cells by drug conditions, we observed clear separation of subtypes (mature CMs’ and ‘immature cells’) based on expression of selected protein markers. We confirmed these subtypes by embedding overlay with independent marker expression patterns (**Fig. 3B, Fig. S2A-B**). That finding highlights that intrinsic biological heterogeneity in our iPSC-CMs dataset dominates over treatment-induced effects. Importantly, approximately 30% of proteins identified as significantly altered between the two subtypes were detected exclusively with AIP.

We observed very striking differences (completely opposite trends in abundance) between the mature CMs and immature cultured cells at their baseline conditions (0µM PR-364, **Fig. 3D**). Immature cells show high expression of proteins associated with epithelial-like phenotype, cell–cell adhesion, and less differentiated status (progenitor-like phenotype), including epithelial intermediate filaments (keratins, KRT19, KRT7, epithelial cells marker, S100A2), desmosome adhesion (plakophilin, PKP1, corneodesmosin, CDSN) and cell growth and differentiation associated proteins (Calmodulin-like5, CALML5, and transporter of epithelial growth factor receptors, ANKS1A). Conversely, mature CMs displayed a functional CM-like proteome, characterized by sarcomere structure components, metabolic competence/maturity, and stress adaptation. Specifically, they show high expression of proteins associated with contractility (MYH6, MYH7, TPM1, ACTA1, FLNC), cytoskeleton remodeling (TAGLN, LASP1, CNN2, ANXA6), metabolism/ mitochondrial function (GAPDH, PKM, VDAC2, ACOT1), and protein folding (heat shock proteins HSPA5, HSP90AA1, HSPA8, HSPB6, HSPA4L).

Importantly, the improved sensitivity of the timsUltra AIP significantly expanded relevant proteome coverage when analyzing the drug effects in subtypes, allowing for more comprehensive pathway analysis. Pathway enrichment of biological effects of PR-364 and per-cell activity inference consistently showed that mature CMs are characterized by elevated oxidative phosphorylation and TCA cycle activity, in agreement with their more developed metabolic phenotype. Immature cells were enriched for pathways related to cytoskeletal structure and organization, as well as developmental processes. PR-364-induced responses were not uniform across the two cell types or biological processes and pathways. Dose-response analyses revealed a non-linear trend in pathway enrichment, i.e., metabolic pathways in mature CMs showed a progressive increase in enrichment with drug concentration, whereas immature cells exhibited more variable responses (**Fig. 3D, Fig. S3**).

At low, intermediate and with maximal activation at the highest dose mature CMs show enrichment in mitochondrial respiration and energy metabolism (*e.g*., TCA cycle, Pyruvate metabolism, Aerobic Glycolysis, Biological oxidation), indicating high-energy metabolic programming, preserved metabolic specialization and stronger oxidative capacity. At intermediate dose (10µM PR-364) mature CMs also shift to suppressing ECM and adhesion related pathways (Cell-cell junction organization, Glycoprotein biosynthetic process, Adheren junction interactions, Vesicle budding, Golgi associated vesicle biogenesis) and shift to glycolipid catabolism at the highest dose (20µM PR-364). Under the lowest dose the immature cells, however, remain more focused on structural remodeling (formation of tubulin folding intermediates, PTMs of tubulin), opening of the cation ion channels (*e.g*., Assembly and cell surface presentation of NMDARs), and most pronounced cytosolic cell respiration (*e.g*., Activation of Ampk, Glycolysis and gluconeogenesis, Aerobic glycolysis), consistent with their less differentiated phenotype. At the intermediate and highest dose immature cells display dominant ECM remodeling (*e.g*., Collagen formation, Extracellular matrix organization, Focal adhesion, Collagen metabolic process, ECM proteoglycans), suggesting stronger interaction with matrix restructuring. Immature cells also show suppression of proliferation and differentiation programs (*e.g*., Intermediate filament organization, Epidermal cell differentiation, Epidermis development) at the highest dose. Comparison with simulated bulk profiles confirmed that averaging across all cells masks important effects associated with specific subtypes.

Several limitations should be considered despite the significantly improved proteome depth; SCP remains incomplete with respect to the cardiac coverage using bulk proteomics. The classification of iPSC-CM subtypes relies on predefined marker sets and may not fully capture the complexity of differentiation or maturity states, which can be examined with pseudo-time and cell trajectory analyses. Furthermore, transcriptomics or functional assays could help redefine cell subtypes. Finally, iPSC-derived cardiomyocytes do not fully recapitulate the phenotype of adult cardiomyocytes. Experiments involving isolated primary cardiomyocytes are required for broader biological understanding of true human cardiomyocyte heterogeneity.

## Conclusions

We demonstrate that integration of the timsUltra AIP into the SCP workflow significantly enhances proteome depth, and reduces data missingness, particularly facilitating detection of low-abundance proteins. These improvements translate directly into enhanced biological insights, enabling high-resolution characterization of iPSC-CMs subtypes and their differential responses to pharmacological perturbation.

## Materials and Methods

### Sample Preparation

The CS0202iCTR (male), CS0083iCTR (female), and CS008iCTR (female) human iPSC lines were generated by the Cedars-Sinai Medical Center iPSC Core from peripheral blood mononuclear cells of a healthy male individual with nonintegrating oriP/EBNA1 plasmids, which allowed for episomal expression of reprogramming factors and shown to be fully pluripotent[30, 31]. iPSCs were maintained in mTESR1 medium on Matrigel-coated cell culture plates and passed every 5 days at split ratios from 1:6 to 1:12 as needed using Versene. Only iPSCs between passage 17 and passage 35 were used for differentiation in this study[31]. iPSCs were differentiated into cardiomyocytes using the established monolayer differentiation protocol utilizing small molecule modulators of Wnt signaling[30]. Differentiated iCMs were metabolically purified by depriving cells of glucose, as previously demonstrated[30]. Purified iPSC-CMs expressed standard cardiac sarcomeric markers cardiac troponin T (cTnT) and α-actinin. Cells were treated with the proteasome inhibitor PR-364 at 0, 3, 10, and 20 µM concentrations (6 µM excluded due to sample preparation issues). Single-cell isolation was performed using the cellenONE to dispense cells into separate wells on a 384-well low binding PCR plate (Biorad HSP3801) containing 200 nl of lysis buffer (100 mM TEAB, 0.2% DDM, 10 ng/nl trypsin). After lysis by freeze-thaw, all samples were subjected to trypsinization with 40 ng/μl trypsin, 4 h incubation at 37 °C, followed by acidification with 0.1% formic acid. All five timepoint 384-well plates were digested in the same batch and frozen until undergoing analysis for single-cell proteomics as outlined in our previously published work [4].

### Mass Spectrometry analysis

Liquid chromatography–mass spectrometry (LC–MS) analyses were performed using a nanoElute 2 system (Bruker Switzerland AG) coupled to either a timsTOF Ultra 2 or the new timsUltra AIP mass spectrometer (Bruker Daltonics GmbH & Co.). Peptide samples (approximately 250 pg to 5 ng in 1 µL) or single cells samples were directly loaded onto a 15 cm C18 reversed-phase column (75 µm inner diameter, 1.7 µm particle size; Aurora Series Elite, IonOpticks). The nanoElute 2 was used with the dissolve sample function method to resolubilize dried down peptide samples from the 384 well plates immediately prior to sample injection. Tryptic peptides were separated at a flow rate of 300 nL min^−1^ using a linear gradient from 5% to 30% buffer B (99.9% acetonitrile, 0.1% formic acid) over 17 min, followed by an increase to 90% buffer B for 3 min, resulting in a total run time of 20 min. The column temperature was maintained at 50 °C using an integrated column heater. DIA was performed using parallel accumulation–serial fragmentation (dia-PASEF). The acquisition method employed 22 isolation windows in 10 MSMS ramps, each 33 Da wide with a 1 Da overlap, and incorporated variable ion mobility settings. Two acquisition steps comprising ten MS/MS ramps were used, resulting in a total cycle time of 1.18 s. The mass-to-charge (m/z) range was set from 100 to 1,700, and the ion mobility range (1/K_0_) spanned 0.7 to 1.30 V·s cm^−2^. Ramp and accumulation times were both set to 100 ms. Collision energy was dynamically adjusted from 20 to 59 eV across the ion mobility range. On the timsUltra 2, the standard collision cell was set radiofrequency to 1500 Vpp, transfer time of 60us and and the pre-pulse storage time was set to 12 µs. On the timsUltra AIP, collision cell radiofrequency was set 1500 Vpp, AIP lens profiles were configured identically for MS1 and MS2 scans, using an alternating current from 800 to 0 Vpp in a slope with a rate of -25 Vpp µs^−1^. Storage-time ramping for MS1 was applied across an ion mobility range of 0.8 to 1.4 1/K_0_ (V·s cm^−2^), corresponding to storage times ranging from 10 to 18 µs. The MS2 storage time was fixed at 17 µs.

### Peptide Identification and Quantification

Raw DIA data were processed using DIA-NN 2.0.2 Academia[32] in library-free mode with the match-between-runs (MBR) algorithm enabled. Precursor and protein-level FDR were controlled at 1%. Quantification was performed at the precursor level, and protein groups were assembled using the MaxLFQ/Quant UMS algorithm[33] as implemented in DIA-NN.

### Quality Control and Preprocessing

Quality control was performed on a per-plate basis. QC metrics extracted from DIA-NN output included the number of precursors and proteins identified per cell, total MS1 and MS2 signal intensity, chromatographic peak width (FWHM RT), median mass accuracy (MS1), normalization instability, and median RT prediction accuracy. Cells failing QC thresholds were excluded from downstream analysis.

Protein detection rate per cell was computed as the fraction of non-zero protein quantifications. Gene name resolution was performed to enable direct gene symbol indexing in the AnnData object. Nine duplicate gene names were resolved: six bovine contaminants (COL3A1, CAP1, ENO1, APOA1, COL1A1, COL1A2) were disambiguated with a “_BOVIN” suffix, two cRAP database contaminants (ALB, LYZ) with species suffixes, and one biological duplicate (TMPO) was split into TMPO_alpha and TMPO_beta isoforms. Seven entries with NaN or empty gene names were recovered from the UniProt Protein.Names column. The original numeric index was preserved in .var[‘original_index’] for back-reference.

### Normalization and Dimensionality Reduction

Raw protein intensities were median normalized per plate to account for differences in total signal across cells. Normalized values were log1p-transformed (log(x + 1)) to stabilize variance. Technical covariates (log-transformed number of detected proteins, normalization instability) were regressed out using Scanpy’s regress_out function[12]. Data were then scaled to unit variance per protein with a maximum clipping value of 10. Principal component analysis (PCA) was computed on the scaled data, retaining 50 components per plate. A shared nearest-neighbor graph was constructed using 15 neighbors in the PCA space, and two-dimensional embeddings were generated using UMAP[34] with default parameters. Community detection was performed using the Leiden algorithm[35] at a resolution of 1.0 (yielding 8 clusters). All clustering used random_state=42 for reproducibility.

### Cardiomyocyte Subtype Classification

CM subtype classification was informed by the reference marker panel from Ai. L *et. al*.[4], which defines mature and structural/immature CM signatures across 13 marker proteins. Initial evaluation of the full 13-marker panel revealed that metabolic markers (*e.g*., GOT1/AATC, MDH1/MDHC, GPI/G6PI) and endoplasmic reticulum markers (*e.g*., PPIB, ANXA5, PDIA3) were pan-expressed across all cells and provided poor discriminatory power. Sub-module analysis identified contractile proteins (*e.g*., MYH7, MYH6) versus cytoskeletal proteins (*e.g*., TPM4, FLNA) as the most informative bimodal markers. A maturity score was computed per cell as the mean log1p expression of myosin heavy chains 7 and 6 (MYH7 and MYH6), and a structural score as the mean log1p expression of TPM4 and FLNA. A CM ratio was defined as the difference between maturity and structural scores. Classification was performed using a 2-component Gaussian Mixture Model (GMM; scikit-learn v1.x [36], GaussianMixture with n_components=2, random_state=42, n_init=10), trained exclusively on 0 µM (vehicle control) cells and an orthogonal 0 µM control set using StandardScaler-normalized maturity and structural scores as features. The 2-component model was selected based on biological motivation (two expected CM populations) rather than BIC, which suggested 4–5 components. The trained model was propagated to all cells via predict and predict_proba [36]. Mean classification confidence (maximum posterior probability) was 0.995–0.996, with zero cells below 0.7 confidence.

### Cross-Validation of CM Subtype Labels

Classification was independently validated using scientist-provided markers: MYH6 as a mature CM marker, and CALD1 (Caldesmon), MYL6, and MYL12B as immature CM markers. A maturity ratio was computed as MYH6 expression minus the mean of CALD1, MYL6, and MYL12B expression (all from the log1p layer). Agreement with the GMM classification was assessed: 85.6% (P3) and 79.0% (P4) of cells classified as Mature_CM had a positive maturity ratio, while only 1.5% (P3) and 0.0% (P4) of cells classified as Immature_CM had a positive ratio. The 15–20% discordance in the Mature_CM group was attributable to MYH6 dropout (51–62% detection rate), with GMM classification sustained by MYH7 detection.

### Differential Expression Analysis

Differential expression (DE) testing was performed within each CM subtype, comparing each PR-364 dose against the 0 µM vehicle control. For P4 dose-response analysis, data were restricted to Female_Line2 cells to avoid cell-line confounding. The Wilcoxon rank-sum test was applied via scanpy.tl.rank_genes_groups (scanpy v1.x;[12]) on the log1p expression layer with tie correction enabled. Multiple testing correction was performed using the Benjamini-Hochberg procedure, and proteins with adjusted p-value < 0.05 were considered significant. A detection rate filter was applied: only proteins detected (non-zero log1p expression) in ≥25% of cells in at least one group (treated or control) were included in testing. Contaminant proteins (bovine serum albumin, cRAP database entries) were excluded from all downstream interpretation.

### Protein Classification: Quantitative vs. Switch-Like

Differentially expressed proteins were classified into three categories based on detection patterns: (1) quantitative - detected in ≥25% of cells in both groups, reflecting graded expression changes; (2) switch-gained - detected in ≥25% of treated cells but <5% of control cells, reflecting drug-induced expression or detection; (3) switch-lost - detected in ≥25% of control cells but <5% of treated cells, reflecting protein loss (consistent with cell death or complete silencing). This classification addresses the missing-not-at-random (MNAR) nature of mass spectrometry missingness[37], in which extreme fold-changes in presence/absence proteins can dominate continuous enrichment analyses.

### Gene Set Enrichment Analysis

Preranked gene set enrichment analysis (GSEA) was performed using gseapy (v1.1.11; [38]) with the prerank function. A composite ranking metric was computed for each protein as sign(log2FC) × −log_10_(p_raw), applied only to quantitative proteins (excluding switch-like and contaminant proteins). This metric balances effect size and statistical significance, avoiding extreme values due to presence/absence artifacts. Rankings were deduplicated by gene symbol, retaining the entry with the highest absolute score. Gene set databases were obtained from MSigDB v2024.1.Hs [39, 40]: Hallmark (50 gene sets), Reactome (1,736 gene sets;[41]), GO Biological Process (7,608 gene sets;[42]), and WikiPathways (830 gene sets; [43]). GMT files were downloaded directly from the Broad Institute MSigDB repository. GSEA parameters: 1,000 permutations, minimum gene set size 10, maximum 500, random seed 42. Pathways with FDR q-value < 0.25 (Hallmark) or < 0.05 (Reactome, GO BP, WikiPathways) were considered significant. Over-representation analysis (ORA) was performed separately on switch-like protein lists using gseapy’s enrich function (Fisher’s exact test) against GO Biological Process gene sets, with all non-contaminant-tested proteins as the background.

### Per-Cell Pathway Activity Scoring

Per-cell pathway activity scores were computed using scanpy.tl.score_genes [44], which calculates the mean expression of pathway gene sets minus the mean expression of a reference set of randomly selected genes with similar expression levels. Pathway gene lists were derived from GSEA lead genes (the leading-edge subset driving enrichment) from the most significant comparison for each pathway, supplemented with curated gene lists for the ubiquitin-proteasome system and intermediate filament/cytoskeleton modules. Pathways scored included: TCA cycle, glycolysis, oxidative phosphorylation/electron transport chain, collagen biosynthesis, ubiquitin-proteasome system, pyruvate metabolism, cytoskeleton/intermediate filaments, and wound healing/fibroblast pathways. Scores were computed on the log1p expression layer.

### Simulated Bulk Comparison

To demonstrate the value of single-cell resolution, a “simulated bulk” analysis was performed by averaging pathway activity scores across all cells within each dose condition (irrespective of CM subtype). This population-level average was compared against subtype-stratified means to visualize how opposing biological responses in Mature CMs and Immature cells are obscured when analyzed at the bulk level.

### Software and Reproducibility

All analyses were performed in Python 3.11 on an HPC cluster (SLURM scheduler) using Jupyter notebooks with interactive, block-by-block execution. Key packages: scanpy [12], pandas, numpy, matplotlib, seaborn, scikit-learn, scipy, statsmodels, gseapy v1.1.11 [38], and openpyxl. Figures were generated at 600 DPI with Arial font for publication compatibility (PDF and PNG formats, with pdf.fonttype=42 for editable text). Intermediate data were saved as h5ad (AnnData) files at each processing stage. Random seeds were fixed at 42 throughout for reproducibility.

## Acknowledgements

Cedars-Sinai Medical Center iPSC Core for human IPSC-lines, Cedars Sinai Proteomics and Metabolomics Core for access to mass spectrometers, and Progenra Inc for providing PR-364.

## Author contributions and conflict of interest

D.A., M.W., M.K., and D.H. are employees of Bruker and may hold financial interests in the company.

## Data availability

Raw data will be available at the PRIDE repository upon manuscript publication.

## Figure Legends

**Supplementary Figure S1.**
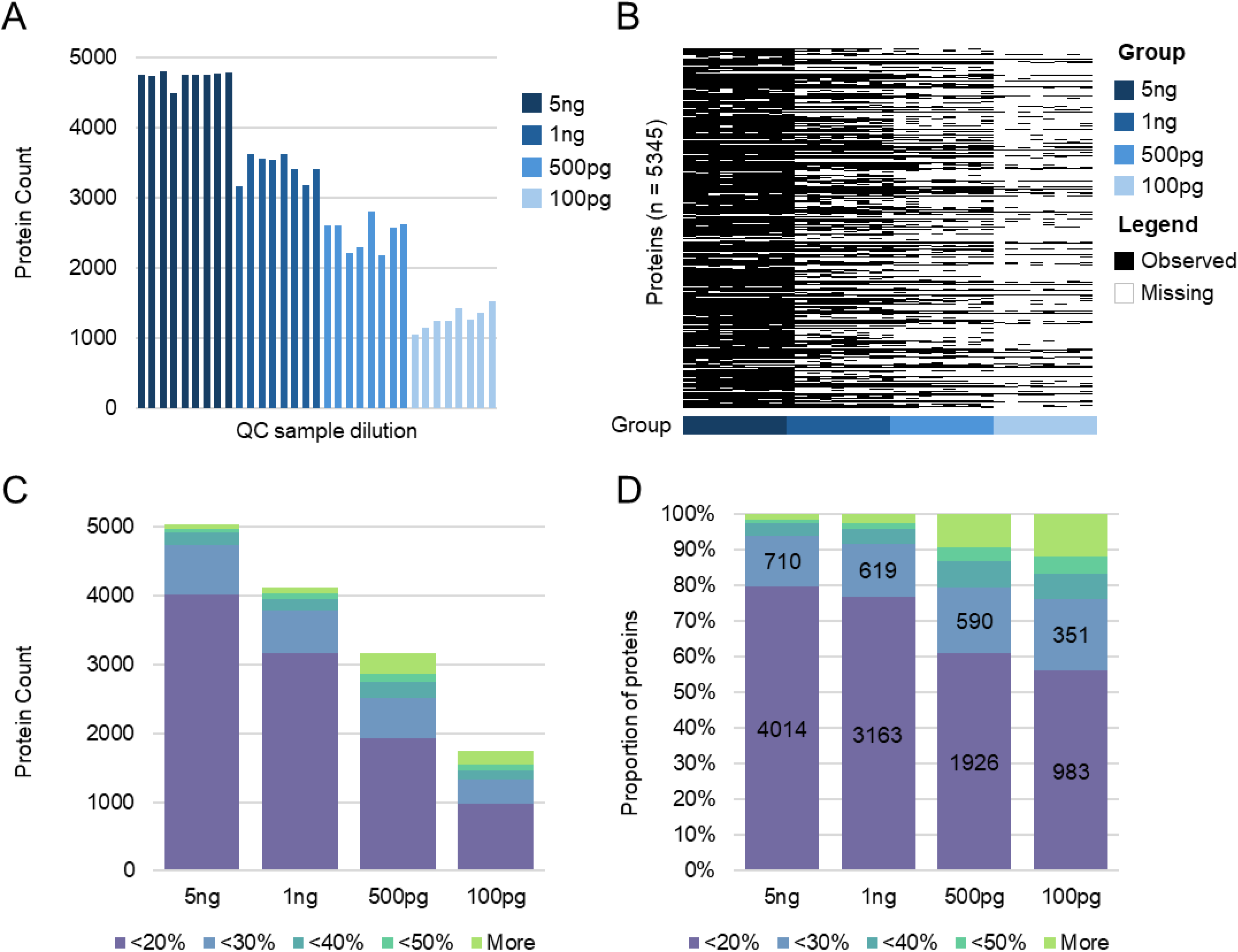
Quality control and quantitative reproducibility across different iPSC-CMs protein lysate concentrations. **A)** Protein counts identified in quality-control samples across decreasing protein amounts (5 ng, 1 ng, 500 pg, and 100 pg). Bar plots show protein identifications per replicate. **B)** Missingness heatmap of proteins detected across different protein inputs. Black indicates observed values and white indicates missing values. Samples are grouped by varying protein amounts (5 ng, 1 ng, 500 pg, and 100 pg). **C)** Absolute number of protein identifications stratified by percent coefficient of variation (%CV) bins across different protein amounts. **D)** Percentage of protein identifications by %CV bins across different protein amounts, illustrating overall good quantitative performance (>75% of all proteins show <20% CV) at 1-5ng with slightly decreased quantitative reproducibility at lower inputs (500pg and 100pg).

**Supplementary Figure S2.**
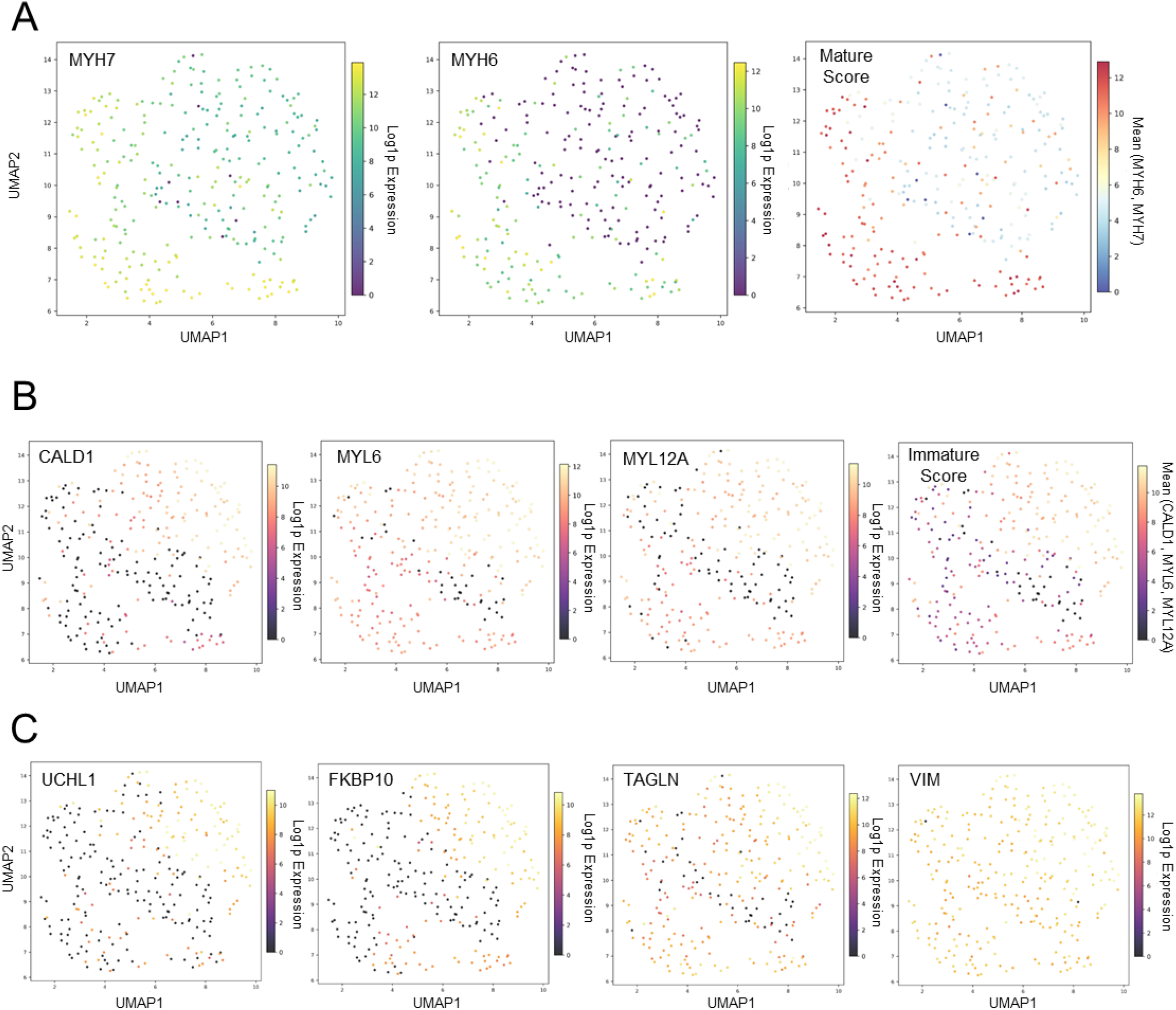
UMAP visualization of cardiomyocyte subtype marker proteins and DEPs distinguishing mature and immature iPSC-CMs subpopulations. **A)** UMAP projections showing expression of previously established (Ai et al. 2025) cardiomyocyte maturity markers. Protein intensity overlays showing MYH7, MYH6, and an aggregated mature cardiomyocyte score, highlighting enrichment of mature cell populations across the UMAP embedding. **B)** UMAP plots of previously established (Ai et al. 2025) protein markers associated with immature cardiomyocytes phenotype. Protein intensity patterns of CALD1, MYL6, and MYL12A, along with an aggregated iPSC-CMs immature score, demonstrate coherent clustering of immature iPSC-CMs populations. **C)** UMAP visualization of selected key DEPs. Protein intensity overlay plots for UCHL1, FKBP1A, TAGLN, and VIM illustrate subtype-specific expression patterns, further validating molecular differences between mature and immature iPSC-CMs states.

**Supplementary Figure S3.**
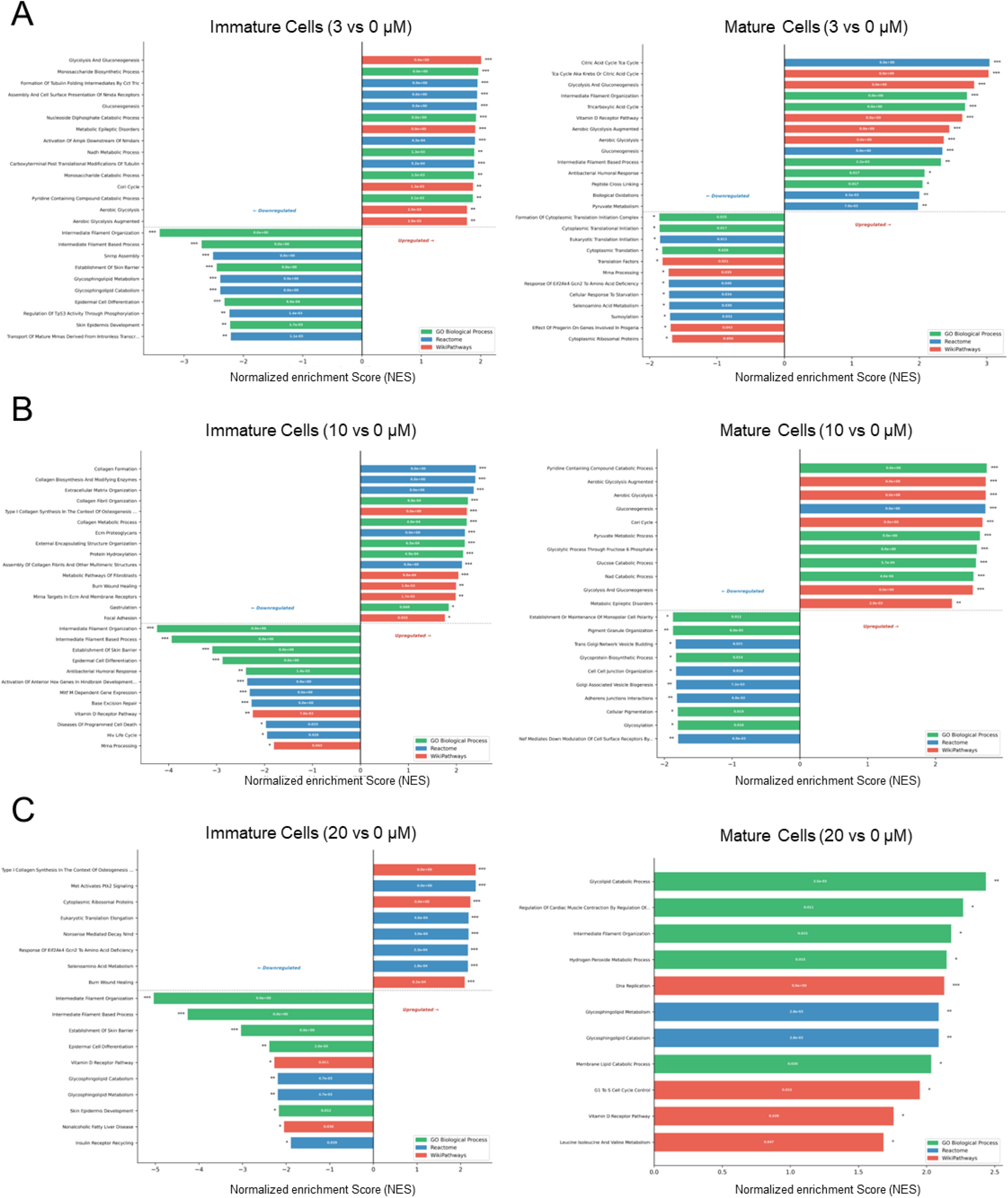
Pathway enrichment analysis reveals dose-dependent functional differences between immature and mature iPSC-CMs. Gene set enrichment analysis (GSEA) of differentially expressed proteins comparing iPSC-CMs at increasing drug concentrations (3 µm, 10 µm, and 20 µm) to the control (0 µm). Left panels show enrichment results for immature iPSC-CMs, and right panels for mature iPSC-CMs. Across all concentrations, immature iPSC-CMs show enrichment of pathways associated with cellular organization, signaling, and developmental processes, whereas mature CMs are enriched for pathways related to energy metabolism, contractile function, and mitochondrial activity. Increasing drug concentrations impact the pathways detection depth and consistency. **A)** Pathway enrichment at 3 µm drug dose. **B)** Pathway enrichment at 10 µm drug dose. **C)** Pathway enrichment at 20 µm drug dose. Bar plots display significantly enriched pathways (FDR < 0.05) from the Gene Ontology Biological Process (GO BP), Reactome, and WikiPathways databases, respectively: green, blue, and red. Positive normalized enrichment scores (NES) indicate pathways upregulated in iPSC-CMs relative to control (concentration 0 µm), while negative NES indicate downregulated pathways.

**Supplementary Figure S4.**
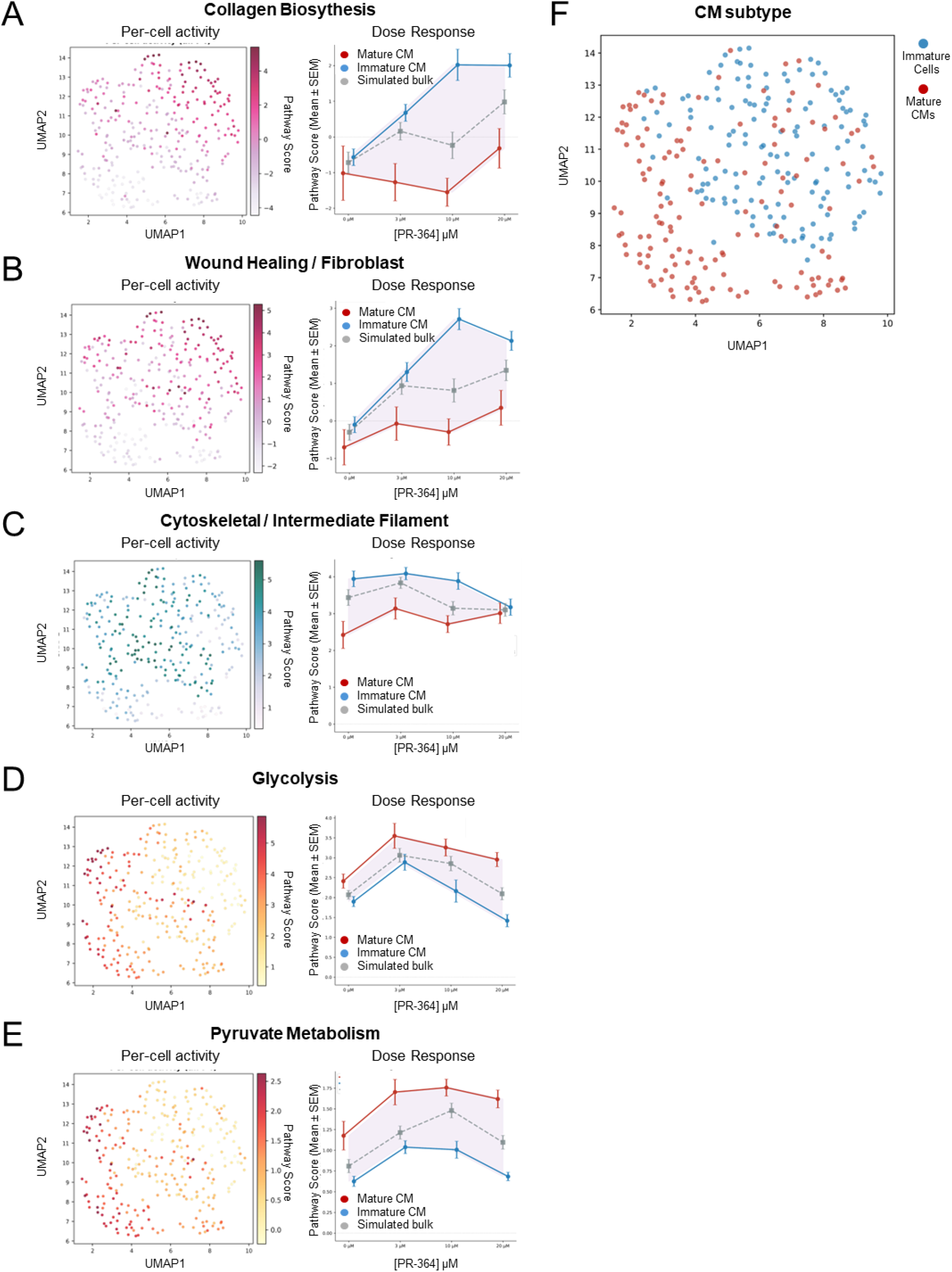
Pathway activity per-cell mapping and dose-response analysis of crucial metabolic functions in iPSC-CMs subtypes. UMAP visualization and dose-response analysis of inferred pathway activities across single iPSC-CMs. For each pathway, the left panel shows per-cell activity scores projected onto the UMAP embedding, the middle panel displays iPSC-CMs subtype using our reference annotation (mature vs. immature), and the right panel presents dose-response curves comparing single-cell measurements to simulated bulk profiles across increasing PR364 drug concentrations. AIP-enabled measurements capture biologically meaningful pathway activity gradients and preserve subtype-specific functional signatures across dose response. **A)** Collagen biosynthesis pathway activity, showing enrichment in fibroblast-like or remodeling-associated single cell states and increasing signal with higher PR364 drug concentrations. **B)** Wound healing activity, highlighting subcluster localization consistent with non-contractile or reparative phenotypes and dose-dependent sensitivity. **C)** Cytoskeleton pathway activity, demonstrating subtype-specific structural organization patterns across UMAP embedding. **D)** Glycolysis pathway activity, enriched in specific iPSC-CMs subpopulations and displaying a non-linear response across input levels. **E)** Pyruvate metabolism pathway activity, reflecting metabolic state differences between mature and immature iPSC-CMs and consistent scaling with increasing PR364 drug concentrations. Dose-response plots (right panels) show mean pathway activity across drug concentrations (3 µm, 10 µm, 20 µm) for mature, immature iPSC-CMs and simulated bulk profiles.

**Supplementary Table 1.** DEPs in SCP PR-364 response study exclusively detected using AIP.

| CM Subtype | Comparison | Total DEPs | AIP Exclusive | Detected Both | Not in AIP Study | % AIP Exclusive | Key AIP Exclusive Proteins |
| --- | --- | --- | --- | --- | --- | --- | --- |
| Mature CM | 3 vs 0 μM | 0 | 0 | 0 | 0 | 0 | |
| Mature CM | 10 vs 0 μM | 0 | 0 | 0 | 0 | 0 | |
| Mature CM | 20 vs 0 μM | 46 | 12 | 4 | 30 | 26.1 | STOM, MACF1, YBX3, COL4A2, PIGS |
| Immature CM | 3 vs 0 μM | 168 | 17 | 143 | 8 | 10.1 | ACTG2, PGK2, TUBA1C, MYL12A;MYL12B, TAGLN2 |
| Immature CM | 10 vs 0 μM | 34 | 11 | 8 | 15 | 32.4 | ALDH1L2, PGM5, PRAF2, CRIP2, CNN1 |
| Immature CM | 20 vs 0 μM | 1514 | 457 | 713 | 344 | 30.2 | CRIP2, FKBP10, FKBP9, P3H1, COL1A2 |
| **ALL (unique)** | **Aggregate** | **1577** | **469** | **736** | **372** | **29.7** |  |

## REFERENCES

1. Brunner, A.D., et al., Ultra-high sensitivity mass spectrometry quantifies single-cell proteome changes upon perturbation. Mol Syst Biol, 2022. 18(3): p. e10798.

2. Hou, W., et al., A statistical framework for differential pseudotime analysis with multiple single-cell RNA-seq samples. Nat Commun, 2023. 14(1): p. 7286.

3. Saddic, L., et al., Single-Cell Proteomics Reveals Novel Cell Phenotypes in Marfan Mouse Aneurysm. Mol Cell Proteomics, 2026. 25(4): p. 101549.

4. Ai, L., et al., Single-Cell Proteomics Reveals Specific Cellular Subtypes in Cardiomyocytes Derived From Human iPSCs and Adult Hearts. Mol Cell Proteomics, 2025. 24(9): p. 100910.

5. Straubhaar, J., et al., Single cell proteomics analysis of drug response shows its potential as a drug discovery platform. Mol Omics, 2024. 20(1): p. 6–18.

6. Petrosius, V., et al., Quantitative Label-Free Single-Cell Proteomics on the Orbitrap Astral MS. Mol Cell Proteomics, 2025. 24(6): p. 100982.

7. Bubis, J.A., et al., Challenging the Astral mass analyzer to quantify up to 5,300 proteins per single cell at unseen accuracy to uncover cellular heterogeneity. Nat Methods, 2025. 22(3): p. 510–519.

8. Ye, Z., et al., Enhanced sensitivity and scalability with a Chip-Tip workflow enables deep single-cell proteomics. Nat Methods, 2025. 22(3): p. 499–509.

9. Ai, L., et al., Enhanced Parkin-mediated mitophagy mitigates adverse left ventricular remodelling after myocardial infarction: role of PR-364. Eur Heart J, 2025. 46(4): p. 380–393.

10. Feyen, D.A.M., et al., Metabolic Maturation Media Improve Physiological Function of Human iPSC-Derived Cardiomyocytes. Cell Rep, 2020. 32(3): p. 107925.

11. Vanderaa, C. and L. Gatto, Revisiting the Thorny Issue of Missing Values in Single-Cell Proteomics. J Proteome Res, 2023. 22(9): p. 2775–2784.

12. Wolf, F.A., P. Angerer, and F.J. Theis, SCANPY: large-scale single-cell gene expression data analysis. Genome Biol, 2018. 19(1): p. 15.

13. Vanderaa, C. and L. Gatto, The Current State of Single-Cell Proteomics Data Analysis. Curr Protoc, 2023. 3(1): p. e658.

14. Schoof, E.M., et al., Quantitative single-cell proteomics as a tool to characterize cellular hierarchies. Nat Commun, 2021. 12(1): p. 3341.

15. Li, A., et al., Giant mitochondria in cardiomyocytes: cellular architecture in health and disease. Basic Res Cardiol, 2023. 118(1): p. 39.

16. Caudal, A., et al., Human induced pluripotent stem cells for studying mitochondrial diseases in the heart. FEBS Lett, 2022. 596(14): p. 1735–1745.

17. Lazar, E., et al., Spatiotemporal gene expression and cellular dynamics of the developing human heart. Nat Genet, 2025. 57(11): p. 2756–2771.

18. Yu, Y., et al., Research trends and hotspots of the applications of single-cell RNA sequencing in cardiovascular diseases: a bibliometric and visualized study. Ann Med Surg (Lond), 2024. 86(12): p. 7164–7177.

19. Deshmukh, R.S., K.A. Kovacs, and A. Dinnyes, Drug discovery models and toxicity testing using embryonic and induced pluripotent stem-cell-derived cardiac and neuronal cells. Stem Cells Int, 2012. 2012: p. 379569.

20. Pang, L., et al., Workshop Report: FDA Workshop on Improving Cardiotoxicity Assessment With Human-Relevant Platforms. Circ Res, 2019. 125(9): p. 855–867.

21. Karakikes, I., et al., Human induced pluripotent stem cell-derived cardiomyocytes: insights into molecular, cellular, and functional phenotypes. Circ Res, 2015. 117(1): p. 80–8.

22. Shahannaz, D.C., et al., Arrhythmogenic Risk in iPSC-Derived Cardiomyocytes: Current Limitations and Therapeutic Perspectives. Medicina (Kaunas), 2025. 61(11).

23. Gherghiceanu, M., et al., Cardiomyocytes derived from human embryonic and induced pluripotent stem cells: comparative ultrastructure. J Cell Mol Med, 2011. 15(11): p. 2539–51.

24. Joshi, J., et al., Human induced pluripotent stem cell-derived cardiomyocytes (iPSC-CMs) for modeling cardiac arrhythmias: strengths, challenges and potential solutions. Front Physiol, 2024. 15: p. 1475152.

25. Simons, E., B. Loeys, and M. Alaerts, iPSC-Derived Cardiomyocytes in Inherited Cardiac Arrhythmias: Pathomechanistic Discovery and Drug Development. Biomedicines, 2023. 11(2).

26. Burnett, S.D., et al., Human induced pluripotent stem cell (iPSC)-derived cardiomyocytes as an in vitro model in toxicology: strengths and weaknesses for hazard identification and risk characterization. Expert Opin Drug Metab Toxicol, 2021. 17(8): p. 887–902.

27. Huang, M., et al., Progress in engineering functional cardiac tissues from iPSC-derived cardiomyocytes: advances in construction and applications. Acta Biomater, 2025. 205: p. 141–163.

28. Clark, A.P., T. Krogh-Madsen, and D.J. Christini, Stem cell-derived cardiomyocyte heterogeneity confounds electrophysiological insights. J Physiol, 2024. 602(20): p. 5155–5162.

29. Churko, J.M., et al., Defining human cardiac transcription factor hierarchies using integrated single-cell heterogeneity analysis. Nat Commun, 2018. 9(1): p. 4906.

30. Sharma, A., et al., Derivation of highly purified cardiomyocytes from human induced pluripotent stem cells using small molecule-modulated differentiation and subsequent glucose starvation. J Vis Exp, 2015(97).

31. Laperle, A.H., et al., iPSC modeling of young-onset Parkinson’s disease reveals a molecular signature of disease and novel therapeutic candidates. Nat Med, 2020. 26(2): p. 289–299.

32. Demichev, V., et al., DIA-NN: neural networks and interference correction enable deep proteome coverage in high throughput. Nat Methods, 2020. 17(1): p. 41–44.

33. Cox, J., et al., Accurate proteome-wide label-free quantification by delayed normalization and maximal peptide ratio extraction, termed MaxLFQ. Mol Cell Proteomics, 2014. 13(9): p. 2513–26.

34. Becht, E., et al., Dimensionality reduction for visualizing single-cell data using UMAP. Nat Biotechnol, 2018.

35. Traag, V.A., L. Waltman, and N.J. van Eck, From Louvain to Leiden: guaranteeing well-connected communities. Sci Rep, 2019. 9(1): p. 5233.

36. Tanaka, T., [[Fundamentals] 5. Python+scikit-learn for Machine Learning in Medical Imaging]. Nihon Hoshasen Gijutsu Gakkai Zasshi, 2023. 79(10): p. 1189–1193.

37. Lazar, C., et al., Accounting for the Multiple Natures of Missing Values in Label-Free Quantitative Proteomics Data Sets to Compare Imputation Strategies. J Proteome Res, 2016. 15(4): p. 1116–25.

38. Fang, Z., X. Liu, and G. Peltz, GSEApy: a comprehensive package for performing gene set enrichment analysis in Python. Bioinformatics, 2023. 39(1).

39. Subramanian, A., et al., Gene set enrichment analysis: a knowledge-based approach for interpreting genome-wide expression profiles. Proc Natl Acad Sci U S A, 2005. 102(43): p. 15545–50.

40. Liberzon, A., et al., The Molecular Signatures Database (MSigDB) hallmark gene set collection. Cell Syst, 2015. 1(6): p. 417–425.

41. Gillespie, M., et al., The reactome pathway knowledgebase 2022. Nucleic Acids Res, 2022. 50(D1): p. D687–D692.

42. Gene Ontology, C., The Gene Ontology resource: enriching a GOld mine. Nucleic Acids Res, 2021. 49(D1): p. D325–D334.

43. Martens, M., et al., WikiPathways: connecting communities. Nucleic Acids Res, 2021. 49(D1): p. D613–D621.

44. Satija, R., et al., Spatial reconstruction of single-cell gene expression data. Nat Biotechnol, 2015. 33(5): p. 495–502.

